# Dual stochasticity in the cortex as a biologically plausible learning with the most efficient coding

**DOI:** 10.1101/811646

**Authors:** Jun-nosuke Teramae

## Abstract

Neurons and synapses in the cerebral cortex behave stochastically. The advantages of such stochastic properties have been proposed in several works, but the relationship and synergy between the stochasticities of neurons and synapses remain largely unexplored. Here, we show that these stochastic features can be inseparably integrated into a simple framework that provides a practical and biologically plausible learning algorithm that consistently accounts for various experimental results, including the most efficient power-law coding of the cortex. The derived algorithm overcomes many of the limitations of conventional learning algorithms of neural networks. As an experimentally testable prediction, we derived the slow retrograde modulation of the excitability of neurons from this algorithm. Because of the simplicity and flexibility of this algorithm, we anticipate that it will be useful in the development of neuromorphic devices and scalable AI chips, and that it will help bridge the gap between neuroscience and machine learning.

## Introduction

Neurons in the cortex continuously generate irregular spike trains with fluctuating membrane potentials and greatly varying firing rates, even across trials in which an animal exhibits appropriate responses and learning in a precisely repeatable manner [1-7]. It has also been found that synapses in the cerebral cortex behave stochastically [8]. The formation, elimination and volume change of dendritic spines exhibit random fluctuations [9-15]. The release of neurotransmitter from synapses is also an inherently stochastic process [16-18].

Theoretically, it has been pointed out that algorithms incorporating these stochastic features can carry out nearly optimal computation in a noisy environment through Bayesian inference [19-28]. To this time, however, the stochastic behaviors of neurons and synapses have been studied separately. It remains unclear if the apparent advantage gained from stochasticity depends on both neurons and synapses behaving stochastically, and if so, whether there is a synergetic interaction between these two types of stochastic behavior. It is also uncertain how learning generated by stochastically functioning neurons and synapses can yield appropriate and precisely repeatable behavioral responses.

In this paper, we show that the stochastic behaviors of neurons and synapses can be inseparably integrated into a simple framework of a sampling-based Bayesian inference model, in which their synergy provides an effective and flexible learning algorithm that is consistent with various experimental findings of the cortex. The derived algorithm accurately describes the plasticity of cortical synapses [29, 30], while it faithfully generates the extremely different timescales of neural and synaptic dynamics, the higher-order statistics of the topology of local cortical circuits [31], and the response properties of cortical neurons, including Gabor-filter-like receptive fields [32, 33], a positive relationship between the receptive field correlation and average connection weight between neurons [34], and the nearly optimal power-law scaling of population activity of neuron [7]. These results strongly suggest that the stochastic behaviors of neurons and synapses are both essential attributes of neural computation and learning. As an experimentally testable prediction of the proposed model, we derive the slow retrograde modulation of the excitability of neurons by postsynaptic neurons. As far as the author is aware, this is the first prediction of its kind and experiments to verify the existence of such slow retrograde modulation have not yet been attempted.

The proposed algorithm can be regarded as a natural integration of the conventional learning theories, error backpropagation learning [35, 36], Bayesian inference [21], Boltzmann machine learning [37], and reinforcement learning [38]. This algorithm can be regarded as an extension of a Boltzmann machine, and it acts as a stochastic variant of error backpropagation. However, the proposed algorithm overcomes most of the limitations that caused backpropagation not to be regarded as the learning principle of the brain. Unlike backpropagation learning, the proposed algorithm does not require neither objective functions, the fine-tuning of parameters, coordinated or synchronous updates of variables, a feed-forward network structure, or the alternating execution of forward and backward computations [35]. Instead, learning is realized through repetition of local and asynchronous stochastic updates of states of neurons and synapses in a network. Because the algorithm is not derived as an optimization of objective functions, it rarely exhibits serious overfitting. We also discuss the close relationship between our algorithm and the temporal difference (TD) learning [38] of reinforcement learning.

## Results

### Neural networks

Most connections between cortical neurons are redundantly realized by multiple synapses [27, 39]. Introducing this redundancy and stochasticity of the neurons and synapses, we model a neural network whose connections are all realized by multiple synapses. Within this model, each neuron and synapse is represented by a binary stochastic variable (Fig. 1a, Methods). The value of a neural variable determines whether that neuron generates a spike, whereas the value of a synaptic variable determines whether that synapse contacts a dendrite of the postsynaptic neuron. A neuron in the network receives inputs from presynaptic neurons and generates a spike with a probability that is a function of the sum of these inputs (see Methods). External inputs, including the target outputs of supervised learning, are presented to the network as variables for some neurons, namely visible neurons, are fixed to values of these input variables. These input data, thus, should be represented by binary vectors. Other neurons in the network, which do not receive external data directly, are referred to as “hidden neurons”.

**Figure 1.**
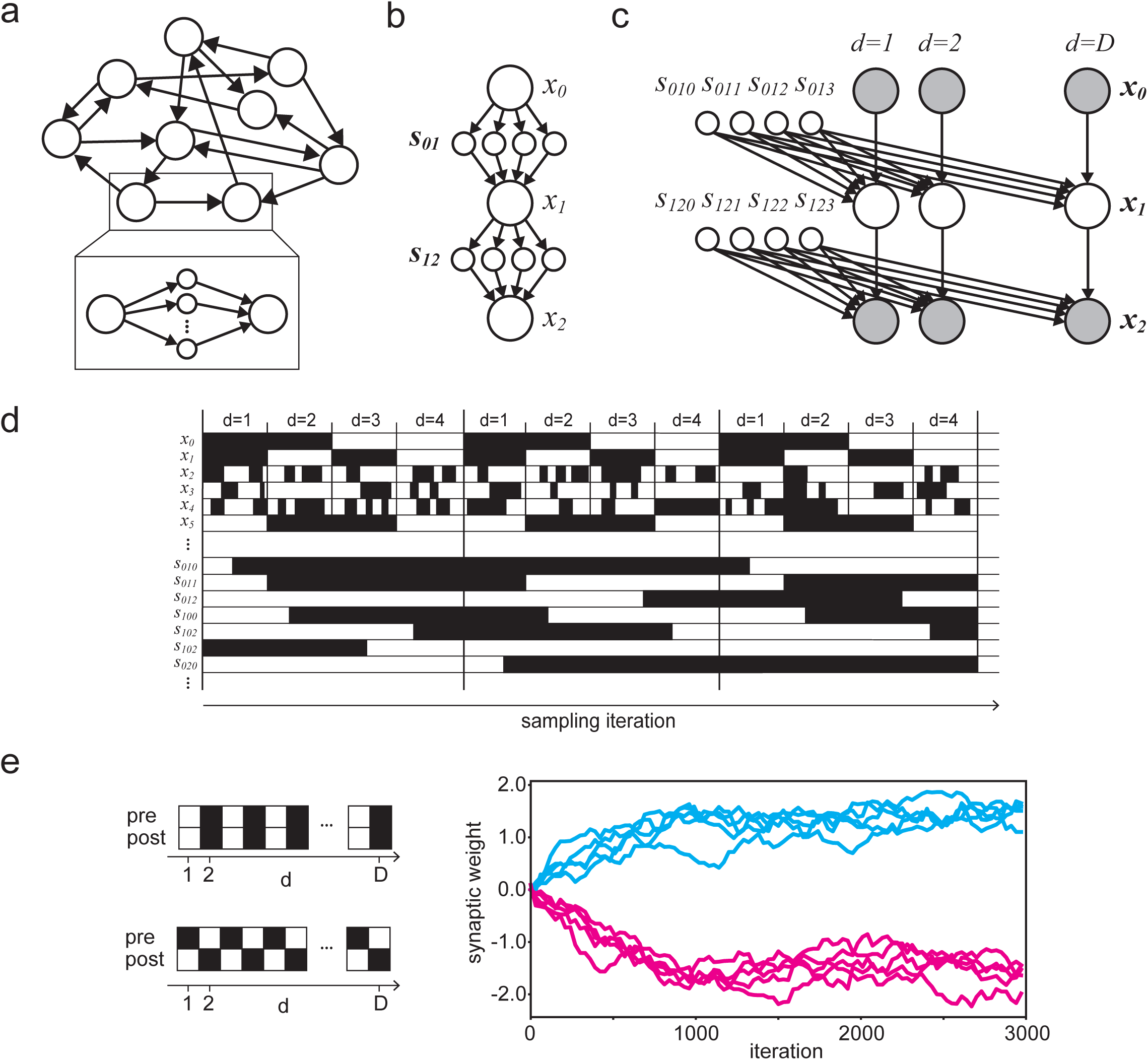
Learning as a Gibbs sampling of synapses and neurons. (a) A neural network is modeled as a population of neurons connected to each other via multiple synapses. (b) A simple neural network consists of three neurons and four synapses per connection. The input, hidden, and output neurons are denoted by *x*_0_, *x*_1_ and *x*_2_, respectively. (c) A graphical model representation of the network shown in (b) in the case that a dataset consisting of D data is presented to the network. Note that the synaptic variables are shared by all data in the dataset while neural variables are not. The white and gray circles represent free and fixed variables, respectively. (d) Schematic of stochastic evolution of neural and synaptic variables. Synapses must evolve much more slowly than neurons to allow the network to incorporate many, ideally all, data in the dataset. Note that values of the visible neurons, *x*_0_, *x*_1_ and *x*_5_ in the figure, are fixed to each datum in the given dataset. (e) Evolution of a synaptic weight (right panel) when presynaptic and postsynaptic neurons fire (left panel) synchronously (cyan) and asynchronously (magenta).

### Biologically plausible learning

With the fundamentals of the network as described above, we formulate learning in the network as a continuing sampling of all the free variables, which include variables representing all synapses and hidden neurons, in the network from a posterior distribution conditioned on the external environment or a given dataset (see Methods). In other words, we hypothesize that the stochastic dynamics of the neurons and synapses in the cortex constitute a continuing random process that aimed at generating a network that suitably interprets the external world.

A sampling from the posterior distribution is computationally and biologically intractable in general, due to the high dimensionality of the system and the complex dependencies among its variables. To solve this problem, we hypothesize that the sampling in the cortex is a Gibbs sampling [40]. The Gibbs sampling ensures that we can replace a sampling from the high dimensional posterior distribution with iterative samplings of each variable from a posterior distribution of that variable conditioned on all other variables. Furthermore, due to the flexibility of the Gibbs sampling, each sampling can be performed in any order and with any frequency. This implies that each neuron and synapse can asynchronously and irregularly update their variables with their own individually determined timings without any global schedule or coordination among them.

Applying Bayes’ theorem to the posterior distributions, we obtain stochastic dynamics, i.e. stochastic update rules, for the neurons and the synapses (see Methods) as

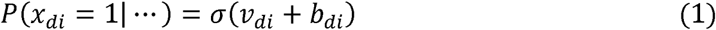

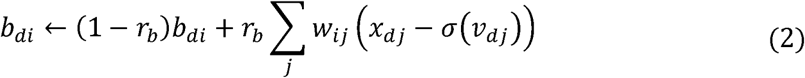

for a neuron and

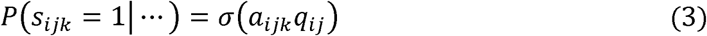

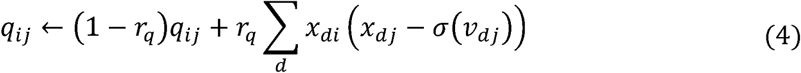

for a synapse, where the dots represent all variables other than the target variable, *x*_*di*_ is the state of the *i*th neuron when the *d*th datum is given to the network, *v*_*di*_ = ∑_*j*_*x*_*j*_*w*_*ij*_ is the membrane potential of the neuron, *s*_*ijk*_ is the state of the *k*th synapse of the connection form *i*th neuron to the *j*th neuron, *r*_*b*_ and *r*_*q*_ are constants which characterize timescales of evolution of *b*_*di*_ and *q*_*ij*_, respectively. *σ(x)*is the sigmoidal function. Following these equations, the state of each neuron and each synapse is repeatedly updated. This simple repetition of irregular and asynchronous stochastic updates is the learning algorithm of the neural network.

Unlike backpropagation learning, in which the alternating execution of forward and backward computations is required, our algorithm realizes learning through the simple iteration of a single computation for each variable. Furthermore, the equations are local in the sense that they depend only on neurons and synapses directly connected to the updated variable, i.e. the Markov blanket of the variable. In addition, the variables do not require any global signals, such as the error of the current output of the network. This asynchronicity and local nature of the dynamics of the learning must be particularly suited to biological implementation of the algorithm.

Note the difference between the summation indices in Eqs. (2) and (4). Synaptic update requires summation over the all data of a given dataset, while neural update does not. This difference is a result of the difference between the data dependencies of the variables (Figure 1b and 1c, see Methods for full details). Because of this difference, synapses need to accumulate the neural activities for many, ideally all, data in the dataset before update their states, while neurons reset their states independently in a manner that depends on each datum individually. This implies that if data are provided sequentially to the network, as in the brain, synapses must evolve much more slowly than neurons (Figure 1d). This explains greatly different timescales of neurons and synapses in the brain. In the following numerical simulations, however, we perform neural updates for all data of the dataset in parallel to accelerates the computational speed.

### Synaptic plasticity

The derived update rule for synapses given in Eqs. (3) and (4) yields plasticity similar to that exhibited by cortical synapses [29]. Each term in the summation in Eq. (3) vanishes unless the presynaptic neuron *x*_*di*_ fires, and if the neuron does fire, whether it will be positive (LTP) or negative (LTD) depends on whether or not the postsynaptic neuron *x*_*dj*_ fires simultaneously. For this reason, each synaptic weight increases if the network receives many data in which the pre-synaptic and post-synaptic neurons of the connection fire synchronously, while it decreases if the pre-synaptic and post-synaptic neurons fire asynchronously (Fig. 1e).

Interestingly, the update rule depends on membrane potential, *v*_*dj*_, in addition to the spikes, *x*_*dj*_, of the postsynaptic neuron. This is consistent with a recently proposed model of STDP that accounts for various properties of synaptic plasticity by introducing the average membrane potential of postsynaptic neurons into the model [30]. It is also noteworthy that, unlike most existing models of synaptic plasticity, the plasticity derived here does not require any artificial bound on synaptic weights. The incremental variation of each synaptic weight automatically decays to zero when the magnitude of the weight becomes very large (Fig. 1e), because *x*_*dj*_ − *σ*(*v*_*dj*_) decays to zero in such cases.

### Retrograde modulation of excitability

The term, *b*_*di*_, of the neural dynamics given in Eq. (2) represents the retrograde modulation of the firing probability of a neuron by its postsynaptic neurons. This term provides a stochastic variant of the error propagation in backpropagation learning. Recall that a target output is given to the network by fixing the states of the corresponding visible neurons to the desired output. Assume that the *j*th neuron is one of these neurons. Then *x*_*dj*_ gives the desired value, and the term *x*_*dj*_ − *σ*(*v*_*dj*_) in the bias *b*_*di*_ of the *i*th neuron gives the difference between the desired value and the expected value of *x*_*dj*_ when the *j*th neuron is not fixed, which is identical to the error in backpropagation learning when the squared error is used as the loss function. Owing to the retrograde modulation, information regarding the desired output provided only to output neurons can spread, i.e. diffuse, over the entire network, even though the variables of the neurons are updated independently, without coordinated scheduling of error backpropagation.

As we will show later, unlike backpropagation learning, the retrograde modulation need not be immediately affected by spikes of the postsynaptic neurons but, rather, can slowly integrate the effects of the spikes. This means that the retrograde bias is determined by the average spike history of the postsynaptic neurons over a finite, presumably quite long, duration. Such slow modulation of the excitability of the neurons could be due to slow changes in the axon initial segment or a long-term modulation of the spike threshold. This seems biologically plausible, as it can be implemented in real cortical circuits. To our knowledge, experiments to verify the existence of such slow retrograde modulation of the excitability of neurons have not yet been attempted.

### Feedforward networks

To understand how the algorithm works, we study its application to a simple problem of supervised learning for a three-layered network (see Methods for full details). Figure 2a displays the evolution of the training and test accuracies as functions of the number of the sampling iteration. It is seen that these accuracies nearly coincide, and they quickly increase to values close to unity and remain there. Significantly, even while these accuracies remain nearly constant, the synaptic weights of the network continue to fluctuate greatly (Fig. 2b), and the firing patterns of the hidden neurons also continue to change, even when the same datum is given to the network, without converging to a fixed pattern (Fig. 2c). These results are consistent with experimental observations of continuing fluctuations of synapses and the trial-to-trial variability of cortical neural activity.

**Figure 2.**
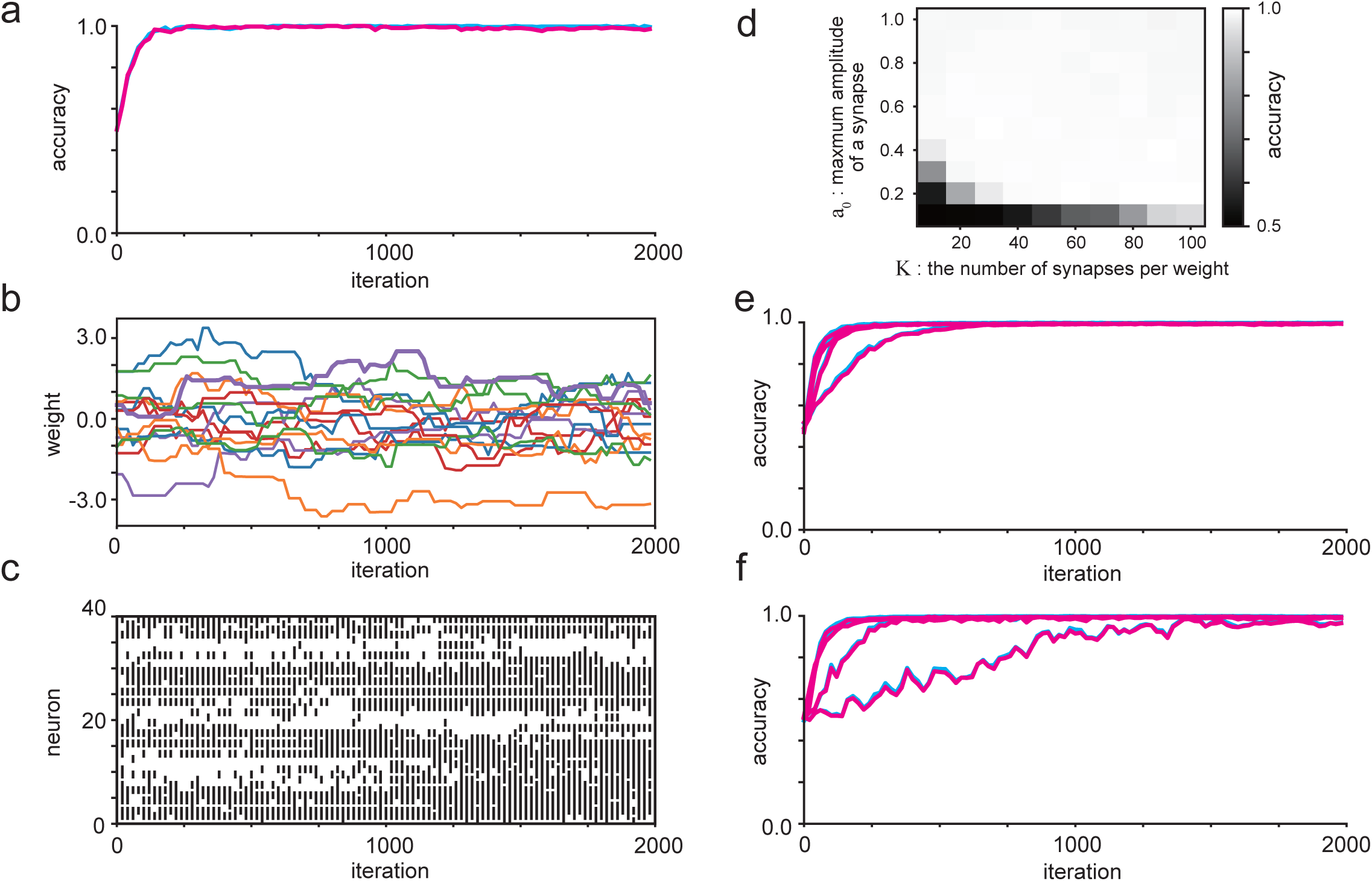
Supervised learning of feed-forward networks using a simple artificial dataset. (a) Training (cyan) and test (magenta) accuracies of a three-layered network as functions of the number of sampling iterations (details given in Methods). (b) Evolution of randomly chosen synaptic weights of the network. Different colors are used for different presynaptic neurons. (c) Evolution of the states of randomly chosen hidden neurons when the learning algorithm with various values of the number of synapses per connection, *K*, the same datum in the dataset is presented to the network. (d) Test accuracies realized by and maximum amplitude of a synapse, *a*_0_. (e) Evolution of accuracies for *r*_*b*_ *=* 1.0, 0.1, 0.01, 0.001, from top to bottom. (f) The same as (i) but for *r*_*q*_ *=* 1.0, 0.1, 0.01, 0.001, from top to bottom.

In order to study the robustness of the algorithm with respect to constants of the algorithm, we calculated the realized test accuracies of the network after learning for various combinations of values of the number of synapses per connection, *K*, and the maximum amplitude of synapses, *a*_0_, (Fig. 2d, see Methos). Except in a narrow range in which one of these constants is so small that the possible maximum weight given by *a*_0_*K* is small, the network almost perfectly learns to perform the task.

Next, we consider the influence of the parameters *r*_*b*_ and *r*_*q*_ (Figure 2e, 2f). Unless *r*_*q*_ is extremely small, and hence *q* evolves extremely slowly, the training and test accuracies will increase and reach values near unity, while their convergence speed decreases as *r*_*b*_ or *r*_*q*_ decreases. (Note that the synaptic evolution is slower than the neuronal evolution even when r_q_ = 1, as discussed above. Therefore, a small value of *r*_*q*_ may result in unrealistically slow synaptic dynamics.) We can thus conclude that the learning is not practically hindered by the use of slow timescales of *b*.

We next study the application of the algorithm to training multilayered feedforward networks using the MNIST dataset to demonstrate the applicability of the method to practical problems (Fig. 3). We found that the accuracies quickly increase to values near 95%, while the number of required iterations and the asymptotically realized accuracies decrease slightly as we increase the number of layers in the network (Fig. 3a-c). Figure 3d displays examples of numerals that the network fails to recognize. These are quite ambiguous and difficult even for a human to identify with confidence.

**Figure 3.**
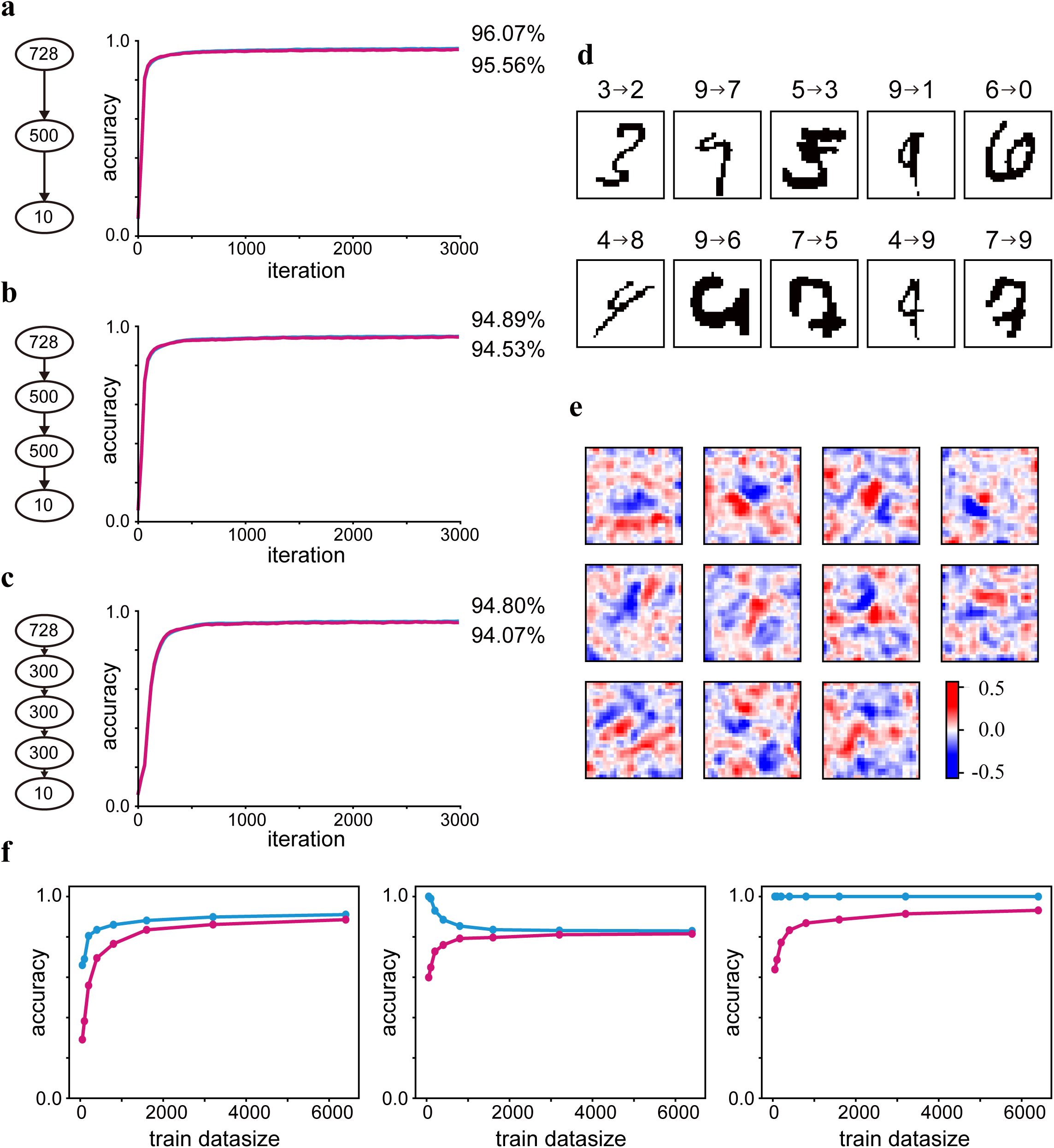
Supervised learning of feed-forward networks using the MNIST dataset. (a-c) Training accuracies (cyan) and test accuracies (magenta) as functions of the number of sampling iterations for (a) three-, (b) four-, and (c) five-layered networks. Each number in a circle indicates the number of neurons in the corresponding layer. (d) Examples of training images (upper row) and test images (lower low) that the three-layered network fails to recognize. (e) Examples of receptive fields of hidden neurons in the three-layered network. (f) Realized accuracies of the three-layered network as functions of the size of the training dataset. The network was trained by the proposed algorithm (left panel), a backpropagation learning algorithm with a naïve stochastic gradient descendent (middle panel), and with the ADAM algorithm (right panel).

Figure 3e illustrates examples of connection weights from the input to the hidden neurons in a three-layered network after training. These correspond to the receptive fields of the hidden neurons. We can see that Gabor-filter-like localized structures that resemble receptive fields of neurons in the primary visual cortex [32] are often organized through the learning. The introduction of a sparse constraint on neural activity into the loss function of learning is known to provide Gabor-filter like receptive fields for an artificial neural network [33]. It is interesting that the network can acquire similar localized structures of receptive fields even without any explicit additional constraint on the learning algorithm.

The learning algorithm can avoid serious overfitting to the training data because it is not derived as a direct optimization of any objective functions. To see this point, we trained a three-layered feedforward network using small numbers of samples of the MNIST dataset, and compared the resulting training and test accuracies with those obtained from backpropagation learning using the stochastic gradient descent (SGD) and ADAM algorithms [41] (Fig. 3f). We found that for both the SGD and ADAM algorithms, as the size of the training dataset is increased, the training accuracy decreases (in the SGD case) or remain nearly constant (in the ADAM case), while the test accuracy increases monotonically, which implies overfitting to the training dataset. Contrastingly, with the proposed algorithm, as the size of the training dataset is increased, both the training accuracy and the test accuracy increase, maintaining a slight difference between them. This implies that the serious overfitting does not occur in the proposed learning algorithm.

### Most efficient power-law coding

A recent experiment carried out by simultaneously recording the activity of a very large number of neurons revealed that the variance spectrum of the principal component of neural activities obeys a power law with an exponent -1.04 that is slightly less than -1 [7]. The authors of the paper [7] proved that if the exponent is greater than -1, the population code by the neurons could not be smooth, while if the exponent is less than -1, high dimensionality of the population code is not fully realized. Thus, the experimentally observed power-law coding with an exponent slightly less than -1 is the most efficient in the sense that in this case, the population response of the neurons lies on a manifold of the highest possible dimension while maintaining high generalizability.

To test whether a network trained by the proposed algorithm realizes the most efficient coding, we numerically calculated the variance spectrum of the principle components of the mean activity of the neurons in the hidden layer of a network trained with the MNIST dataset. As shown in Fig. 4a, the variance spectrum exhibits clear power-law decay with an exponent of -1.06. This is very close to the experimental result, and is indeed slightly less than -1. We conclude that the learning algorithm leads the network to the most efficient coding.

**Figure 4.**
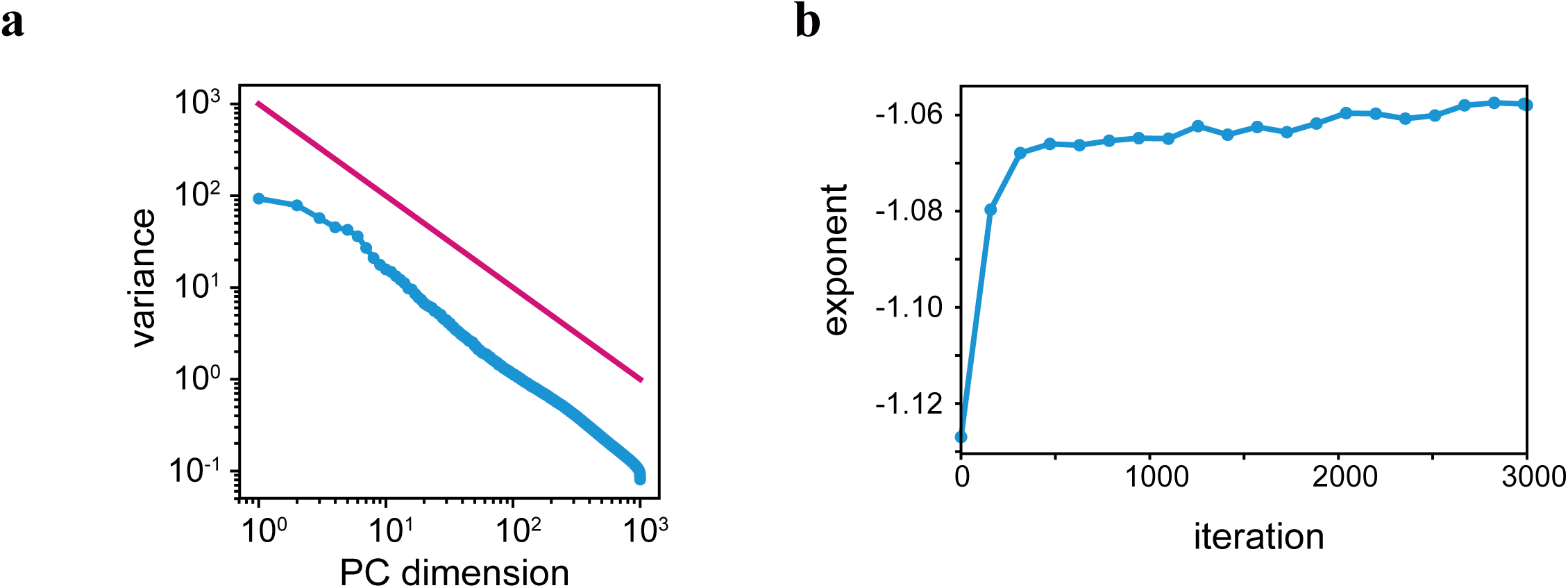
Nearly optimal power-law decay of the variance spectrum for the principal component of the neural activity. (a) Variance spectrum of the principle components of the mean activity of the hidden neurons of the three-layered network after training (cyan). The variances are arranged in descending order. The line (magenta) indicates the critical slope corresponding to an exponent of -1. (b) Evolution of the power-law exponent of the variance spectrum as a function of the number of sampling iterations.

We next study how the exponent of the power law develops during learning. Figure 4b shows that the exponent approaches a value close to -1 from below as the learning proceeds. This result implies that the network first learns a coarse representation of the dataset and then gradually acquires finer structures while maintaining generalizability of the representation of the data in coding space. This leads us to conclude that the robustness or generalizability of the population coding takes priority over the precision of the data representation in the learning. This priority must particularly be beneficial for animals that must survive in a ceaselessly changing environment.

### Recurrent networks

We next applied the algorithm to train a network with recurrent connections (Fig. 5a) using the MNIST dataset. Figure 5b displays the evolution of the training and test accuracies as functions of the number of sampling iterations. The accuracies were obtained from the states of the output neurons of the network measured after recursive evolutions of the states of the hidden neurons. We see that, as in the case of the results for the feedforward networks, the accuracies nearly coincide and rapidly increase. This implies that the algorithm is even able to train a recurrent network and rarely overfits.

**Figure 5.**
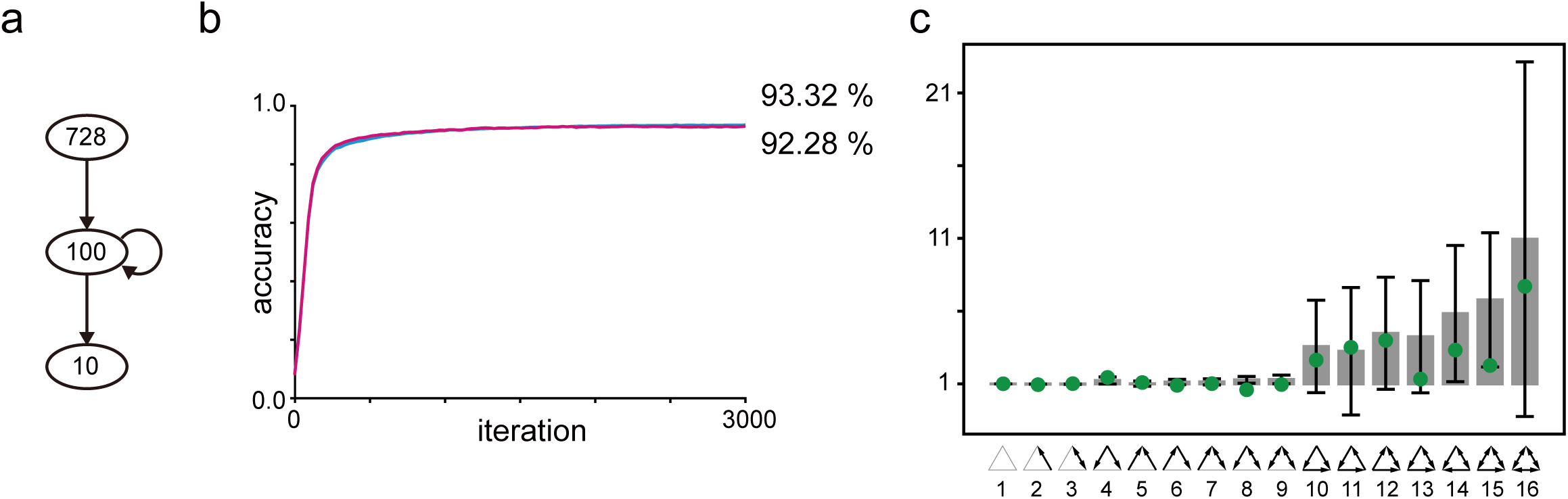
Supervised learning of a recurrent network using the MNIST dataset. (a) A recurrent neural network. Each number in a circle indicates the number of neurons in the corresponding layer. (b) Evolution of the training accuracies (cyan) and test accuracies (magenta) of the recurrent network. (c) Average ratios of the actual numbers of three-neuron patterns in the trained recurrent network to those predicted by the null hypothesis (gray bars), (details given in Methods). The error bars indicate the range of ±2*σ*. The green circles are experimental results for real cortical circuit [31].

### Statistics of network motifs

It has been reported that local cortical circuits are highly nonrandom, and that connectivity patterns consisting of multiple neurons, known as network motifs, exhibit a characteristic distribution in which highly clustered patterns are overrepresented [31]. To study whether a recurrent network trained by the proposed algorithm acquires a similar distribution of connectivity patterns, we determined connectivity of triplets of neurons in a trained recurrent network. The statistics for the ratio of the actual counts of triplet patterns to the chance level are plotted in Figure 5c. The same ratios for the experimental results are also overlaid in the figure. While there are some exceptional cases in which the ratios obtained here are somewhat larger than those obtained experimentally, the two distributions of triplet patterns are surprisingly similar. As observed experimentally, highly-connected motifs, i.e., those numbered 10 through 16 in Fig. 5c, are overrepresented by a factor several times greater than chance level. These results support the validity of the derived algorithm as a model describing the formation of local cortical circuits.

### Connection weights and receptive field correlation

A recent experiment of the primary visual cortex revealed that the connection weights between pairs of pyramidal neurons become stronger as the receptive fields become more similar [34]. To test whether the trained recurrent network accounts for this relationship, we measured the connection weights between pairs of neurons and the receptive field correlations between these neurons (fig. 6a). We found that the average connection weight between neurons is positively correlated with the correlation between the receptive fields of these neurons (Fig. 6b). Particularly, by restricting our analysis to only connection weights with positive values (i.e., *w*_*ij*_ > 0), we were able to reproduce a nonlinear relationship between the average connection weight and the receptive field correlation (Fig. 6c), which was similar with the experimental result [34].

**Figure 6.**
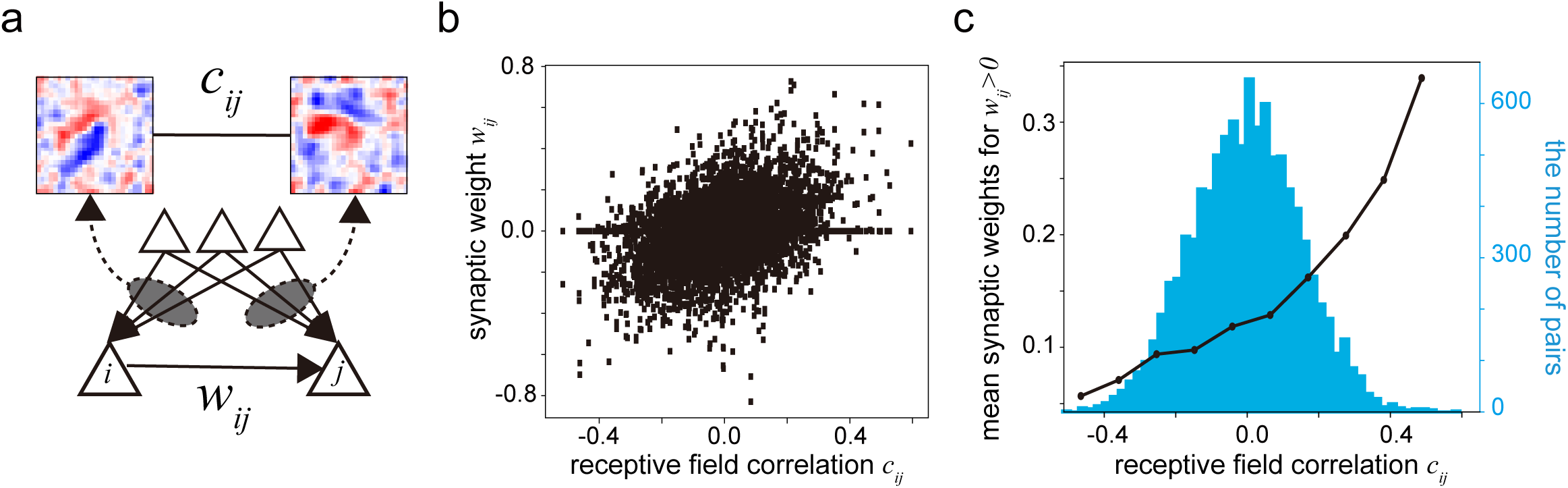
Correlation between connection weights between pairs of neurons and the similarity of the receptive fields of these neurons. (a) Correlation coefficients, *c*_*ij*_, of the receptive fields between pairs of neurons and connection weights, *w*_*ij*_, between these neurons are measured for the recurrent network after training. (b) The connection weights between neurons. (c) The average weight of these connections with positive weights, *w*_*ij*_ > 0, as pairs of neurons positively correlates with the receptive field correlations between these a function of the receptive field correlation. Underlying histogram shows the distribution of the receptive field correlations for the pairs of neurons. These results correspond to Figure 2g and 2i of [34].

### Temporal sequence learning

We next consider the application of the algorithm to train recurrent networks with temporal sequences (Fig. 7). We prepared periodic temporal sequences in which the same temporal inputs may appear multiple times at different times, and trained networks to predict the next input of the current sequence. In this case, networks need to learn to store the history of inputs over some interval to generate the desired output. The training procedure was the same as that used in the case considered in Figs. 2-5, except that we identified the iteration of the updates of the variables of the network as the time development. In contrast to the algorithm known as “backpropagation through time”, this procedure does not require virtually unfolding the recurrent connections of the network along the time axis.

**Figure 7.**
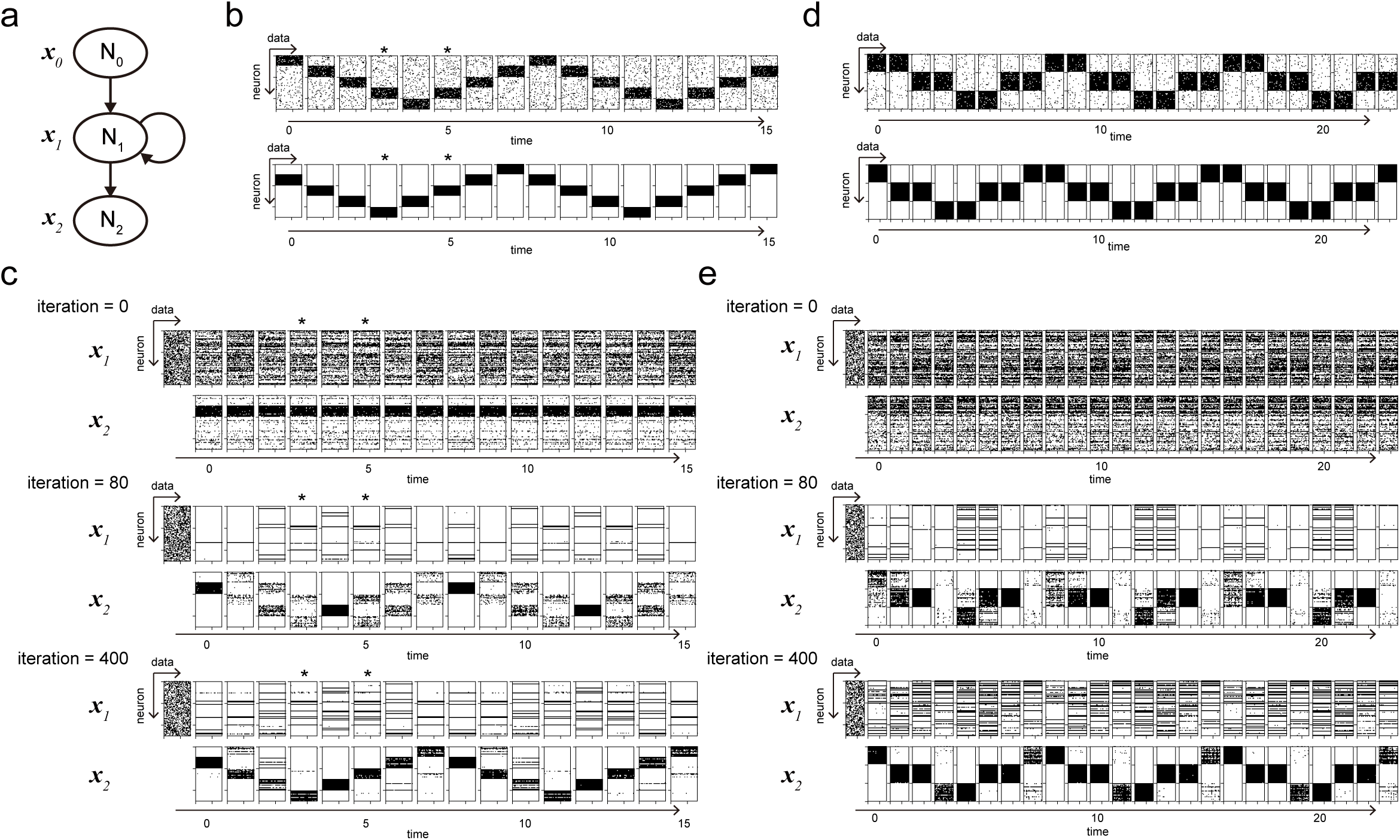
Supervised learning of recurrent networks using temporal sequences. (a) Network structure. (b) Input sequences (upper panels) and target output sequences (lower panels) of the first dataset. The asterisks indicate examples of times at which the network must output different patterns while the current input to the network is the same. (c) Activities of hidden neurons (upper panels) and output neurons (lower panels) after 0, 80, and 400 sampling iterations. (d) Input sequences (upper panels) and target output sequences (lower panels) of the second dataset. (e) The same as (c), but for the second dataset.

We prepared two sequences that require one-step and two-step memories, respectively. For both tasks, the networks were successfully trained to output the desired sequences by the learning algorithm. We find that after learning, the output produced by the network in any given case depends not only on the current input but also on past inputs. This indicates that, through the learning, the algorithm causes the network to store input histories into the activity of the hidden neurons.

## Discussion

In this study, we showed that Gibbs sampling from a joint posterior distribution of neurons and synapses in a network conditioned on an external environment or given dataset explains the stochastic natures of synaptic development and neural activities of cortical circuits. This provides a practical and biologically plausible learning algorithm that yields results consistent with various experimental findings for cortical circuits. The derived stochastic dynamics of synapses are consistent with the plasticity of cortical synapses, and those of neurons naturally describe highly irregular features and the trial-to-trial variability of spike trains of cortical neurons.

The evolution equation for neurons has a term that results in the retrograde modulation of the excitability of a neuron by its postsynaptic neurons. Due to this term, the algorithm acts as a stochastic variant of algorithms with error backpropagation through which target outputs provided to a part of network can spread over the entire network. However, in contrast to the case of backpropagation learning, the retrograde modulation seen in our model requires neither synchronous nor precisely coordinated operations, which are major reason that backpropagation has not been regarded as the learning principle of the brain. Retrograde modulation need not be immediately affected by postsynaptic action potentials but, rather, can slowly integrate postsynaptic spikes. The mechanisms regarded as likely to be responsible for this behavior include the combination of action potential backpropagation from soma to dendritic spines [42] and retrograde transsynaptic transport of certain chemicals [43], glia-mediated modulation [44], and disynaptic connections from postsynaptic to presynaptic neurons. Experimental confirmation of this modulation would be convincing evidence to support the validity of the proposed framework as a learning and computational principle of the brain.

The reason why both neurons and synapses must be stochastic in the cortex can be understood as follows. Suppose that the primary purpose of the cortex is to generate synaptic weights that are consistent with the external environment. Mathematically, this can be formulated by generating samples of synaptic weights from a Bayesian posterior distribution conditioned on the external environment, which is derived from the likelihood function of the synaptic states determined by the dataset. However, when the network has hidden neurons, the likelihood is given by a marginal distribution of the joint conditional distribution of hidden neurons and synapses in which neural and synaptic variables are strongly coupled. Thus, sampling of synaptic states inevitably also requires sampling of neural states. In this way, the stochastic behaviors of neurons and synapses are integrated in the cortical dynamics.

It has been pointed out that the temporal difference (TD) method of reinforcement learning provides a model of spike-timing-dependent Hebbian plasticity [38]. In TD learning, the value function of the current state is updated after the delivery of a reward such that the value function includes the TD error defined as the difference between the value of the delivered reward and its predicted value. Interestingly, each term in the evolution equations of the latent variables of the synapses and neurons, i.e. *b*_*di*_ and *q*_*ij*_, can be regarded as a representation of the TD error of the delivered spikes instead of the rewards. Indeed, each term in Equations (2) and (4) takes the form of a scaled difference between the measured spike of a postsynaptic neuron, *x*_*dj*_, and its expected value, *σ*(*v*_*dj*_), before observation of the spike. In this analogy, therefore, each spike plays the role of a reward for the presynaptic neurons and synapses from these neurons. At the same time, each spike is also transmitted to postsynaptic neurons, thereby changing their firing probabilities in a manner that depends on the corresponding synaptic weights. Thus, each spike in the cortex plays two roles, rewarding backward units and selecting next states by triggering the next spikes. It will be a fascinating topic of future research to theoretically elucidate the relationship between our algorithm and TD learning.

The algorithm developed here is closely related to the Boltzmann machine [37], which is a stochastic recurrent neural network in which neurons are represented by stochastic binary variables and iteratively updated to realize a thermal equilibrium state of a globally defined energy function specified by a given temperature. In that algorithm, symmetric connection weights between neurons are trained with a gradient descent method to make the equilibrium state approximate a target distribution in which the neural states of given a dataset have high probability. The algorithm developed in the work can be regarded as an extension of the Boltzmann machine in which the Bayesian posterior distribution is directly considered, instead of a thermal equilibrium of an energy function, and in which synapses in addition to neurons are modeled as stochastic variables to be sampled. This extension results in biologically realistic update rules of variables that are implemented concurrently for neurons and synapses with no need for explicit switching of different computations or fine scheduling of parameters, such as simulated annealing.

In the limit of a large number of training data, the posterior distribution of Bayesian inference converges to a delta function whose peak position coincides with the result of the maximum likelihood estimation (see Methods). Therefore, if a sufficiently large amount of external data is provided to the network, our learning algorithm almost surely generates synaptic and neural variables that most suitably reflect the external data. This situation contrasts with that for backpropagation learning, in which convergence to an optimal solution is not always guaranteed, even when a large amount of data is used.

Because it seems to operate in accordance with a basic principle of neural computation and learning, we believe that our model provides a theoretical foundation for various experimental findings regarding cortical dynamics and various methods of machine learning. Most of the recent experimental findings regarding neuroscience have not yet been fully utilized in the development of machine learning. This may be because backpropagation learning is not consistent with the functioning of the brain. These experimental results include results regarding short-term plasticity of synapses [45], Dale’s principle, the long-tail distribution of synaptic weights [31], columnar structure, laminar organization, canonical circuits [46, 47], and innate structure formed through developmental stages [48]. Incorporating these features into our model with the goal of clarifying their functional roles is an important future project. Introducing precise continuous dynamics of membrane potentials and spike generation mechanisms into the model are also important topic to be investigated.

## Methods

### Neural networks

Introducing the redundancy and stochasticity of the neurons and synapses, we model a neural network that consists of N neurons connected via K synapses per connection. The connection weight from the *i*th to the *j*th neuron is given as a weighted sum of the synaptic states as 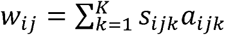, where *s*_*ijk*_ ∈ {0,1} is a binary random variable describing the state of the *k*th synapse of the connection from the *i*th to the *j*th neuron. The weights are generally asymmetric, and we set *w*_*ii*_ = 0 to avoid self-connections. The strength, or amplitude, of a synapse, denoted by *a*_*ijk*_, is a constant that represents the contribution of the synapse to the weight, which can be interpreted as corresponding to the amplitude of the miniature postsynaptic potential (PSP) of the synaptic contact. Note that *a*_*ijk*_ is a constant, and it is fixed during learning. For simplicity, we used an evenly spaced sequence from −*a*_0_ to *a*_0_ for the values of the amplitude throughout the work: *a*_*ijk*_ *= a*_*k*_ *=* (2(*k* − 1) /*k* − 1) *a*_0_ (*k* = 1,2,…, *K*).

The *i*th neuron receives inputs from its presynaptic neurons and randomly generates a spike with the probability *P*(*x*_*i*_ = 1) = *σ*(*v*_*i*_) = *σ*(∑_*j*_ *x*_*j*_ *w*_*ij*_), where the state of the neuron, *x*_*i*_ ∈ {0,1}, is a random binary variable representing the spike firing of the neuron, and *σ*(*x*) is the activation function of a neuron, for which we use the sigmoidal function *σ*(*x*) = (1+ *e*^−*x*^)^−*x*^ throughout the paper. The weighted sum of inputs to the neuron *v*_*i*_ corresponds to the membrane potential of the neuron.

Neurons in the network are classified into three groups, input, output, and hidden. Input and output neurons together are referred to as visible neurons. External data, including the target outputs of supervised learning, are input into the network by fixing the states of visible neurons to the values of the data. Each datum, therefore, must be a binary vector. When we obtain the output of the network after and during learning, we fix only input neurons, keeping output and hidden neurons free.

The states of the neurons, including the hidden and visible neurons, are updated in response to each datum in the dataset as it is input, while the state of a synapse depends on the dataset as a whole, because the aim of the learning is generally to obtain networks, i.e., sets of synaptic states, that consistently reflect all of the data in the dataset (Figure 1c). For this reason, we write the state of the *i*th neuron at the time that the network is receiving the *d*th datum of the dataset as *x*_*di*_, using the data index, while the state of the synapse, *s*_*ijk*_, does not have a data index. Figure 1c presents a graphical model representation of the data dependency of the variables for the simplest case of a three-layered neural network.

### Learning algorithm

The learning of the network is modeled as a Gibbs sampling of all free variables from their posterior joint distribution conditioned on the fixed variables, 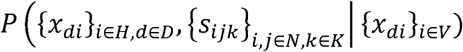. Here, *d* is the data index of the given dataset, *D*, while *V* and *H* denote the sets of visible and hidden neurons, *N* represents the set of all neurons, and *K* denotes the set of synapses for each connection. (The number of variables of the network is thus *ND* + *WK*, where *W* is the number of connections in the network.) The Gibbs sampling allows us to replace sampling from a generally high-dimensional joint distribution with a repetition of samplings of each single variable from a posterior distribution conditioned on all other variables,

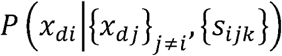

and

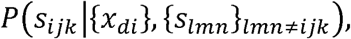

for a neuron and a synapse respectively. In the Gibbs sampling, the order of the samplings need not be fixed, but can be random. Also, the sampling frequencies of different variables can be different. Therefore, in general, the state of each neuron or synapse will change at times that are determined independently for each, depending only on the conditions experienced individually by that neuron or synapse.

To derive an explicit description of the posterior distribution of a neuron, let us consider the log likelihood ratio for *x*_*di*_. Using the Bayes rule, we obtain

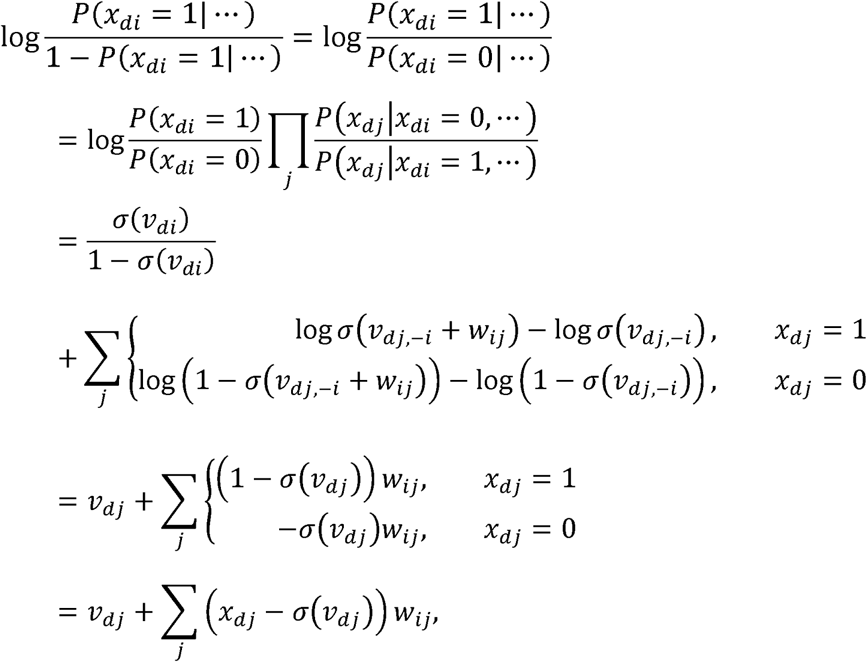

where the dots represent all variables other than *x*_*di*_, i.e., 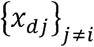 and {*s*_*ijk*_}, and we have *v*_*dj*,−*i*_ = ∑_*k*≠*i*_ *x*_*dk*_*w*_*kj*_. To obtain the 5th line, we have assumed that *v*_*dj*, −*i*_ ≫ *w*_*ij*_ and linearized each term of the summation with respect to *w*_*ij*_. Solving the above equation for *p*(*x*_*di*_ = 1| …), we obtain Eq. (1) in the main text with

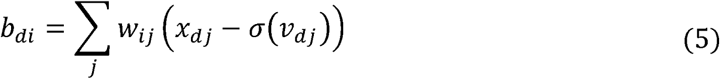

as the posterior distribution (i.e. the stochastic update rule) of the neuron.

Similarly, the log likelihood ratio for the synapse *s*_*ijk*_ is

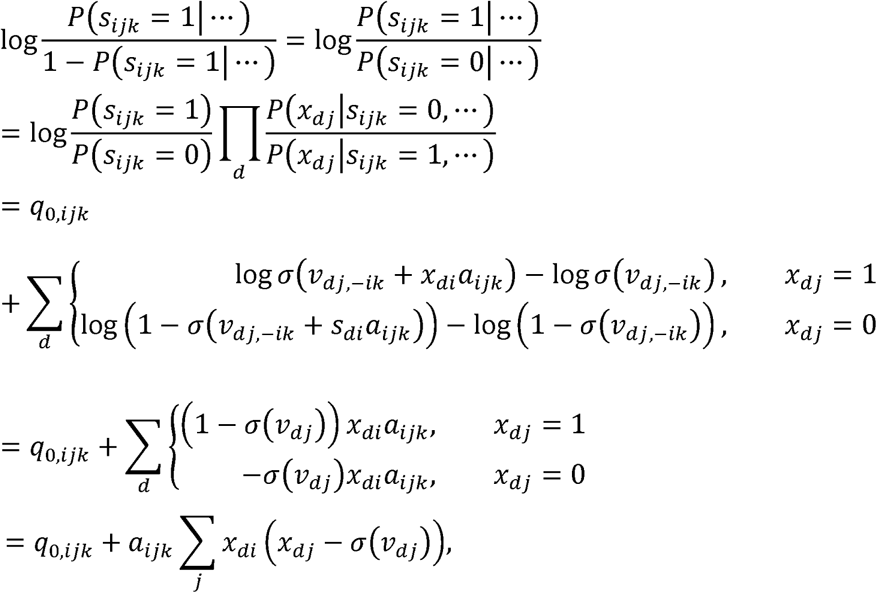

where the dots represent {*x*_*di*_} and {*s*_*lmn*_}_*lmn*≠*ijk*_, and we have *v*_*dj*,−*ik*_ = ∑_*l*_*x*_*l*_*w*_*lj*_ −*x*_*di*_*s*_*ijk*_*a*_*ijk*_. To obtain the 5th line, we have assumed *v*_*dj*, −*ik*_ ≫ *a*_*ijk*_, and approximated each term in the summation as a quantity linear in *a*_*ijk*_. The constant *q*_0,*ijk*_ represents the log likelihood ratio of the prior distribution, *p*(*s*_*ijk*_), which simply vanishes unless the prior distribution is biased. We assumed *q*_0,*ijk*_ = 0 throughout the work. Solving the above equation for *p*(*s*_*ijk*_ = 1| …) gives Eq. (3) in the main text and

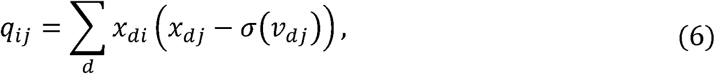

which constitute the explicit description of the posterior distribution or the update rule of the synapse.

Equations (5) implies that a spike of a neuron immediately changes *b*_*di*_ and excitability of presynaptic neurons. However, such immediate retrograde modulation has not been experimentally reported, and seems biologically implausible. Rather, it is biologically more natural that *b*_*di*_ evolves slowly while accumulating the effects of postsynaptic spikes as Eq. (3) in the main text, where *r*_*b*_ characterizes the timescale of the evolution. (Here, *r*_*b*_ satisfies 0 < *r*_*b*_ ≤ 1, while in the case *r*_*b*_ = 1, Eq. (3) reproduces to Eq. (5).) Thus, *b*_*di*_ is determined by the average spike history of the postsynaptic neurons over a finite, presumably quite long, duration. Similarly, we can also generalize the evolution equation for *q*_*ij*_ as Eq. (4) in the main text. As demonstrated in Figure 2e and 2f, these generalizations rarely decrease the learning accuracy of the algorithm.

The following is a possible biological implementation of our algorithm. (i) Each neuron in the network continuously evaluates its membrane potential, *v*, and bias, *b*, and stochastically generates spikes with the probability given in Eq. (1). (ii) A generated spike is immediately integrated into the membrane potentials of its postsynaptic neurons, while it slowly modulates the excitability of its presynaptic neurons (Eq. (2)). (iii) The spike firing also modulates the latent synaptic variable, *q* (Eq. (4)). (iv) The state of each synapse is changed asynchronously and irregularly in accordance with Eq. (3) with a frequency that is sufficiently slow that sensory neurons receive a large variety of external inputs during the average interval of the updates.

### Numerical simulations

All numerical simulations are written in Python, with the open-source matrix library CuPy. In details of the procedures to train a feedforward network are as follows. (i) We first prepare *ND* binary variables for the *N* neurons and *WK* synapses where *D* is the number of data in the training dataset, *W* is the number of connections in the network, and *K* is the number of synapses per connection. (ii) Then we fix the variables of the visible neurons to the values of the data in the training dataset, and initialize the values of the hidden neurons and synapses randomly to 0 or 1 with probability 1/2. (iii) To avoid perfectly synchronized updates, we randomly choose the ratio *r*_*x*_ for the hidden neurons and update their variables according to Eqs. (1) and (2). (iv) Similarly, we randomly choose the ratio *r*_*s*_ for the *WK* synapses to update in accordance with Eqs. (3) and (4). (iv) We repeat (iii) and (iv) as many times as desired. The procedure to obtain the prediction of the network is the same as that for the training procedure, except that we fix only the input neurons and update the hidden and output neurons in accordance with Eqs. (1) and (2), keeping the synaptic values fixed. In order to accelerate the computation, we can use the average activities of the neurons, *σ*(*v*_*di*_), instead of their binary variables, *x*_*di*_, and omit the biases, *b*_*di*_, during the prediction procedure. The training and test accuracies are defined as the ratio of the number of inputs that enables the network to generate the correct outputs to the total numbers of inputs of the training and test datasets.

### Dataset

Except in the cases described by Figs. 2 and 7, we used the MNIST dataset, which consists of a training dataset of 60,000 examples and a test dataset of 10,000 examples in which each image has 28 × 28 pixels. Because pixels in the MNIST data range from 0 to 255, we replaced them with 0 or 1, depending on whether the value of the pixel is below or above 255/2. We thus obtain 784-dimmensional binary input vectors.

In the situation considered in Fig. 2, we trained a three-layered network consisting of 40 input, 40 hidden, and 2 output neurons to learn a simple task that is a noisy and high-dimensional variant of the XOR problem. The datasets were artificially generated as follows. We first prepared two-dimensional binary vectors (*X*_*d*1_, *X*_*d*2_), where *d* is the data index of the dataset. Then, to obtain 40-dimensional binary input vectors (*Y*_*d*1_,…, *Y*_*d*40_), we set *Y*_*di*_ = *X*_*d*1_ for *i* = 1, …, 20 and *Y*_*di*_ = *X*_*d*2_ for *i* = 21, …, 40, and then flipped their values randomly with a probability of 0.1 to obtain randomized input dataset. The desired outputs of the two-dimensional vectors are given by *Z*_*di*_ = (0,1) if XOR(*X*_*d*1_, *X*_*d*2_) = 0 and (0, 1) if XOR(*X*_*d*1_, *X*_*d*2_) = 1. The training dataset and test dataset each contains of 400 examples.

In the situation considered in Fig. 7, we used datasets consisting of temporal sequences to train recurrent networks. Let us write the input data and desired outputs at time t as *X*_*di*_ (*t*) and *Z*_*di*_ (*t*). These are fed into the network by fixing the neurons in the input layer as *x*_0,*di*_ (*t*) = *X*_*di*_ (*t*), 0 < *i* ≤ *N*_0_, and those in the output layer as *x*_2,*di*_ (*t*) = *Z*_*di*_ (*t*), 0 < *i* ≤ *N*_2_, where *N*_0_ and *N*_2_ are numbers of neurons in the input and output layers, respectively. A dataset is prepared as follows. We first prepare an integer sequence *s*(*t*), where 1 ≤ *s*(*t*) ≤ *S* and 1 ≤ *t* ≤ *T*. We then set *X*_*i*_(*t*) = 1 if (*s*(*t*) − 1)*N*_0_/*S* < *i* ≤ *s*(*t*)*N*_0_/*S* and *X*_*i*_(*t*) = 0 otherwise. The target output is set as *Z*_*di*_ (*t*) = *X*_*i*_ (*t*+ 1) if 1 ≤ *t* < *T* and *Z*_*di*_ (*T*) = *X*_*i*_ (0) otherwise. To obtain randomized input vectors *X*_*di*_ (*t*), we replicated *X*_*i*_(*t*) and randomly flipped them as *X*_*di*_ (*t*) = *X*_*i*_ (*t*) with probability 0.9 and *X*_*di*_(*t*) = 1 − *X*_*i*_(*t*) with probability 0.05. The integer sequence *s*(*t*) given by 1, 2, 3, 4, 5, 4, 3, 2, 1, 2, 3, 4, 5, 4, 3, 2 with *N*_0_ = 250, *S* = 5 and *T* = 16 was used as the first dataset and 1, 1, 2, 2, 3, 3, 2, 2, 1, 1, 2, 2, 3, 3, 2, 2, 1, 1, 2, 2, 3, 3, 2, 2 with *N*_0_ = 150, *S* = 3 and *T* = 24 is used for the second dataset.

### Parameters

We used *r*_*q*_ = 1.0, *r*_*s*_ = 0.001, *K* = 200, *a*_0_ = 0.1, and *D* = 100 in the situation considered in Fig. 1e, and *r*_*b*_ = *r*_*q*_ = *r*_*x*_ = 1.0, *r*_*s*_ = 0.001, *m*_0_ = *m*_1_ = 10, *K* = 50, *a*_0_ = 0.5, and *D* = 400 in the situation considered in Fig. 2. In the situation considered in the remaining figures, except Figs. 4, 5, 6 and 7, we used *r*_*b*_ = 0.01, *r*_*x*_ = 0.9, *r*_*q*_ = 0.1, *r*_*s*_ = 0.001, *m*_0_ = *m*_1_ = 20, *K* = 100, and *a*_0_ = 0.1. In the case of Fig. 4, we use *K* = 200, and in the case of Figs. 5 and 6, we used *r*_*x*_ = 0.5, and in the case of Fig. 7, we used *r*_*b*_ = 0.1, *r*_*x*_ = 1, *r*_*q*_ = 0.01, r_s_ = 0.01 and *D* = 4000. The numbers of hidden neurons in the three-layered network that are not specified in the figures are 1000 for Figs. 3f and 4, 100 for Figs. 5 and 6, and 500 for Fig. 7. Connection probability was 1.0 and 0.5 for feed forward connections and recurrent connections, respectively.

## Data analysis

### Spectrum variance of principle components

After we trained the three-layered neural network considered in Fig. 3a using the MNIST dataset, we fixed the synapses and obtained the average activities of the hidden neurons {*σ*(*v*_*di*_)}_*i*∈*H*_. Note that the quantities *v*_*di*_ for the hidden neurons were deterministic in this case because both the input neurons and the synaptic connections from them were fixed. The principle component analysis was applied to the average activities after they were standardized. Then we obtained the explained variance of each principle component, which is the eigenvalue of the covariance matrix of the standardized average activities, and ordered them in descending order. The exponent of the power law was estimated with a least-square linear fit of the variance spectrum in log-log space.

### Statistics of network motifs

We trained a three-layered recurrent network with 100 hidden neurons. Then, we determined the number of connection patterns among the triplets of neurons over all possible combinations of 3 neurons chosen from 100, i.e. for 100 · 99 · 98/6 = 161,700 triples. Here, we only counted connections whose synaptic weights were greater than or equal to 0.27, in order to exclude small and negative connections. Null hypothesis of the counts is defined as the same way that provided in the paper [31]. Namely, we determined the numbers of unidirectional and bidirectional connections in all pairs of neurons and calculated the predicted number of three-neuron patterns by assuming all constituent pairs of neurons in each triplet pattern are chosen independently, while maintaining the probabilities of the measured unidirectional and bidirectional connections. We performed 20 learning trials in order to obtain the mean and standard deviation, *σ*, of the ratio of the actual number of each triplet pattern to that obtained with the null hypothesis.

### Limit of large training data size

Let us represent all free variables of a network consisting of neurons {*x*_*di*_} and synapses {*s*_*ijk*_} collectively by *θ.* We can reasonably assume that the prior distribution *P*(*θ*) is positive for all values of *θ* and that each training datum is independently generated from a data distribution *P*_*d*_ (*x*). Then, the posterior distribution satisfies

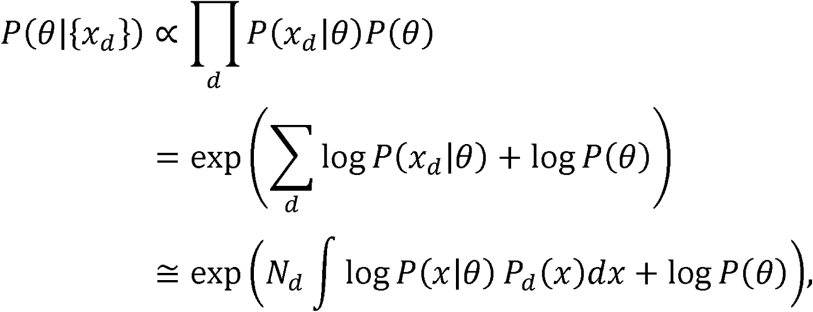

which generally converges to a delta function *δ*(*θ* − *θ*_0_) in the limit of a large number of training data, *N*_*d*_ → ∞, where *θ*_0_ = argmax_*θ*_ ∫ log *P*(*x*|*θ*) *P*_*d*_(*x*)*dx*. (If maximum is realized of multiple values of *θ* simultaneously, the posterior distribution will converge to the sum of the corresponding delta functions.)

